# Integrated cyto-physiological and proteomic analyses reveal new insight into CMS mechanism in a novel upland cotton CMS line LD6A

**DOI:** 10.1101/2022.01.09.475591

**Authors:** Zheng Jie, Aziz Khan, Zhou Bujin, Zhou Qiong, Najeeb Ullah, Kong Xiangjun, Liu Yiding, Liu Fang, Zhou Ruiyang

## Abstract

Cytoplasmic male sterile (CMS) system has extensively been exploited for hybrid vigor in plant breeding programs. However, its application in many crops is limited due to poor understanding of molecular mechanism of fertility restoration. Using advanced analytical approaches, we elucidated molecular pathways regulating CMS induction and fertility restoration in cotton. Reproductive structures of a novel CMS (LD6A) and its maintainer (LD6B) line were analyzed for physiological and proteomic changes during the development process. Significant differential expression of proteins, such as Abrin, malate dehydrogenase, malic enzyme, isocitrate dehydrogenase, histone acetyltransferase was observed in CMS and its maintainer line. Transmission electron micrographs of anther tapetum showed that inner ridge of CMS mitochondria was relatively indistinct than that of LD6B with narrower membranous space at tetrad stage. Further, relatively higher reactive oxygen species were accumulated in the anther of CMS than its maintainer line at pollen mother cell and tetrad stage. We suggest that abnormal sequence of mitochondrial ribosome gene rps4 and rpl10 and high expression of ribosome-inactivating protein gene Abrin in CMS line damaged mitochondrial membrane and consequently induced pollen sterility. These data provide new insight into CMS mechanism in cotton crops and a tool to develop new CMS germplasm resources.

## 1. Introduction

Cotton is a major industrial crop cultivated for natural fiber. Lint yield and fiber quality is strongly regulated by heterosis. In breeding programs, cytoplasmic male sterile (CMS) lines greatly accelerated the exploitation of heterosis in crops, such as rice (Song *et al*., 2021), rape (Wolko *et al*., 2019), wheat (Gupta *et al*., 2019). Since the discovery of first CMS gene *T-urf13* in maize (Levings, 1976), molecular mechanisms of many CMS genes have been improved (Dewey *et al*., 1987; Moneger F, 1994; Song & Hedgcoth, 1994; H *et al*., 1995). For example, in rice wild abortion CMS line, the CMS gene *wa352* was identified after rice mitochondrial genome was sequenced completely(Hu *et al*., 2014). This gene triggers premature tapetal programmed cell death and consequent pollen abortion by interacting with a nuclear coded mitochondrial protein COX11 (Luo *et al*., 2013). Similarly, a CMS protein ORF288 is suggested to induced toxicity to reproductive structures in a rape CMS line (Jing *et al*., 2012). However, molecular mechanisms of CMS induction in cotton plants are still to be explored.

Reactive oxygen species (ROS) are small molecular compounds, which play a key role in growth, development, and defense pathways(Sies & Jones, 2020). These ROS can also induce cytoplasmic male sterility by damaging mitochondrial membrane (Møller, 2001). For example, during microspore development in rice HL-CMS line Yuetai A, an abnormal increase of ROS led to ATP and NADH depletion, degrading mitochondrial genomic DNA (Wan *et al*., 2007). Furthermore, ATP and NADH depletion can block ascorbic acid - glutathione cycle in mitochondria, inducing mitochondrial dysfunction and expression of mitochondrial chimeric open reading frame (Ling Huang, 2011). In cotton, CMS-D8 line Zhong 41A (Yang *et al*., 2018a), Kenaf CMS line H722A (Zhou *et al*., 2019) and P9SA (Liu *et al*., 2021), suppression of ROS elimination ability led to rapid release of ROS and abnormal microspore development.

Biological energy conversion is carried out in mitochondria, especially in membrane protein complex and inner membrane cristea (Kühlbrandt, 2015). Mitochondria contain an independent genome and, gene expression system, such as mitochondrial ribosomes (mitoribosomes) (Tomal *et al*., 2019). The mitoribosomes are mainly composed of mitochondrial ribosomal proteins, which are encoded by nuclear genomic DNAs and rRNAs encoded by the mitochondrion (Robles & Quesada, 2017). Deficient mitochondrial translation could reduce mitochondrial respiratory activity through abnormal membrane structure (Pfeffer *et al*., 2015a) and influence stability and function of mitoribosomes.

The application of next generation sequence (NGS) technology, RNA-seq including transcriptome sequencing and small RNA sequencing are popular in cotton CMS research (Suzuki *et al*., 2013; Yang *et al*., 2014; Wu *et al*., 2017; Kong *et al*., 2017a; Yang *et al*., 2018b; Zheng *et al*., 2019; Shahzad *et al*., 2020; Li *et al*., 2021). Even regulatory pathways of some key genes associated CMS induction were also screened out. However, due to interference of nuclear gene expression and RNA splicing in cotton, CMS induction mechanism in this species is still poorly understood. To better explore CMS mechanism, proteomics approach especially isobaric tags for relative and absolute quantification (iTRAQ) could be an ideal tool. Some special proteins and their potential interaction neatwork related to pollen development and CMS were screened out. In rice HL-CMS line, four key enzymes involved in TCA (tricarboxylic acid) cycle reduced the ATP supply (Sun *et al*., 2009). These include triosephosphate isomerase (TIM), fructokinase II, pyruvate kinase, pyruvate dehydrogenase (PDH). In cotton GMS line 2074A, *GhA11G1250* encoded a mitochondrial localization of peroxisomal-like protein HSP14.5, participated in response to ROS is critical to pollen abortion(Nie *et al*., 2020). UDP-glucuronosyl/UDP-glucosyltransferase, 60S ribosomal protein L13a-4-like, and glutathione S-transferase, as well as heat shock protein Hsp20, ATPase, F0 complex, and subunit D were related to the microspore abortion in Yamian A, a CMS line with a wild cotton genetic background (Zhao *et al*., 2020).

In this study, combined with cytological, physiological, and proteomic analyses, the key regulators of another development were discussed. Through iTRAQ and bioinformatic analyses, differentially accumulated proteins especially in CMS line and its maintainer line will be detected. These proteins will be analyzed as CMS candidate factors, such as toxic protein Abrin, TCA related enzymes MDH, ME, IDH, mitoribosome protein gene *rps4, rpl10*, ROS elimination enzyme SOD, POD, APX. This will help to better understand CMS mechanism in cotton and broaden CMS germplasm resources.

## 2. Materials and methods

### 2.1 Plant Materials

A cotton CMS line LD6A and its maintainer LD6B were planted in the Guangxi University experimental farm (Nanning, April-October) and the National Wild Cotton Nursery (Sanya, October-March) under normal field management conditions. Floral buds during different developmental stages (2-3mm, pollen mother cell stage, 3-4 mm tetrad stage, 4-5 mm early uninucleate, 5-6 mm late uninucleate, >6 mm mature pollen stage) were collected for TEM observation. Pollen abortion stage (tetrad stage, 3-4 mm in diameter) buds were frozen in liquid nitrogen and stored at −80°C for RNA and protein isolation.

### 2.2 Morphological and Cytological Observations

In the full bloom stage, typical flowers of CMS and maintainer line were observed by a digital camera (Canon, Tokyo, Japan) and the photos of flower buds were taken by a stereomicroscope (Olympus, Tokyo, Japan).

The series size flower buds were soaked in Carnoy’s fixation solution for 24 hours, then washed with gradient alcohol. After the fixation, the buds were stored in 75% alcohol. Tissue blocks were dehydrated in ascending series of ethanol, cleared in a series xylene and embedded in paraffin. Then the buds embedded in paraffin were cut into 10 μm pieces sections. The paraffin sections were treated with xylene dewaxing, gradient ethanol hydration, vanadium iron hematoxylin stain, then gradient alcohol and xylene dehydration. Finally, the sectioned floral buds were observed using an electron microscope (Olympus BX53, Tokyo, Japan).

The observation of subcellular microstructure of both lines were detected by TEM (HITACHI H-7650, Tokyo, Japan) at the Guangxi Medical University. The series size flower buds were treated in 2.5% glutaraldehyde fixation solution for 12-48 hours, then 1% OsO_4_ at least 12h under 4 □ condition. After dehydration, infiltration the buds were cut into ultrathin section. Finally, the ultrathin sections were double stained with lead citrate (C_6_H_5_O_7_Pb) and 2% uranium acetate (UO_2_ (Ch_3_COO_2_)_2_), the sections were observed, selected, and photographed under transmission electron microscope (Hitachi H-7650).

### 2.3 Physiological traits related to ROS metabolism

Evaluation for some enzymes related to ROS metabolism in different size flower buds were carried out. Antioxidant enzymes such as aseorbateperoxidase (APX), superoxide dismutase (SOD) and peroxidase (POD) and malondialdehyde (MDA) a marker of plasma membrane peroxidation were quantified (Daniele *et al*., 2005). The standard procedure we followed was described by Sofo et al (Sofo *et al*., 2004). Three biological replicates and three technical replicates were used in each experiment.

### 2.4 Protein extraction and iTRAQ analysis

1~2 grams of floral bud (Tetrad stage) from both LD6A and LD6B with 10% PVPP were grounded into powder in liquid nitrogen and then sonicated on ice in Lysis buffer (8 M Urea and 40 mM Tris-HCl containing 1 mM PMSF, 2 mM EDTA and 10 mM DTT, pH 8.5). After centrifugation, and precipitation, the dried, resuspended samples were incubated for reduction and alkylated. Protein quantitation was detected by Bradford assay and SDS-PAGE. After sample digestion, peptides were labeled (iTRAQ labeling) and peptide fractionation was performed with HPLC.

### 2.5 Bioinformation analysis

Quality control (QC) is performed to determine if a re-analysis step is needed. An automated software IQuant (Wen *et al*., 2014), was applied to the quantification of proteins with a Picked protein FDR strategy (PSM-level FDR≤0.01, Protein-level FDR≤0.01) (Savitski *et al*., 2015). All proteins with a false discovery rate (FDR≤0.01) proceed with downstream analysis including GO, COG and Pathway. Further, we also perform deep analysis based on differentially expressed proteins, including Gene Ontology (GO) enrichment analysis, KEGG pathway enrichment analysis and cluster analysis.

### 2.6 RNA extraction and qRT-PCR validation

Total RNA of both selected lines was extracted from the floral bud (Tetrad stage) using an RNA Isolation Kit (TransGen Biotech, Beijing, China). The relative expressions of DEGs were verified by qRT-PCR and analyzed by 2^-ΔΔCt^ method(Livak & Schmittgen, 2001), where 18s gene was considered as an endogenous control. All the primers were designed by Primer Premier 5.0 and synthesized by BGI (Shenzhen, China). RT-PCR and qRT-PCR were performed according to the previously used method (Kong *et al*., 2017b). Three biological replicates and three technical replicates were used in each analysis.

## 3. Results

### 3.1 Comparative observation of microspore development in morphological characteristics, paraffin section and mitochondrial structure by transmission electron microscope

Morphological variations between LD6A and LD6B flowers were observed across different developmental stages (Figure 1). From pollen mother cell stage to late uninucleate stage, there were no noticeable structural differences in petals, stigma, sepals, and stipules. However, at MP stage, LD6A has a more prominent stigma than LD6B. At the flowering stage, anthers of LD6A were shriveled while that of LD6B were plump, and granular pollen can be observed clearly.

**Figure 1.**
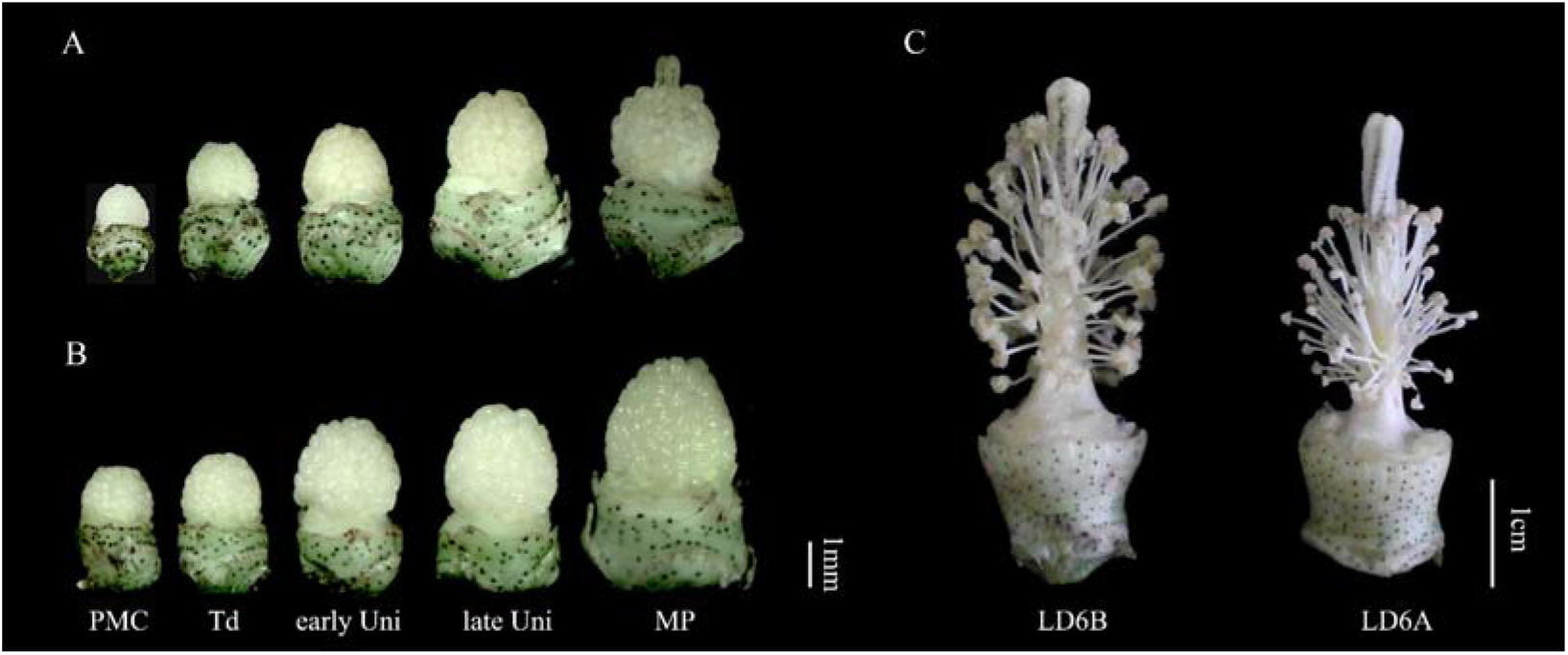
Phenotype of the floral buds of the cytoplasmic male sterility (CMS) line LD6A and its maintainer LD6B: (A): CMS line LD6A; (B): Maintainer line LD6B; (C): blossoming flowers of LD6A and LD6B PMC, pollen mother cell stage; Td, tetrad stage; early Uni, early uninucleate stage; late Uni, late uninucleate stage; MP, mature pollen stage.

**Figure 2.**
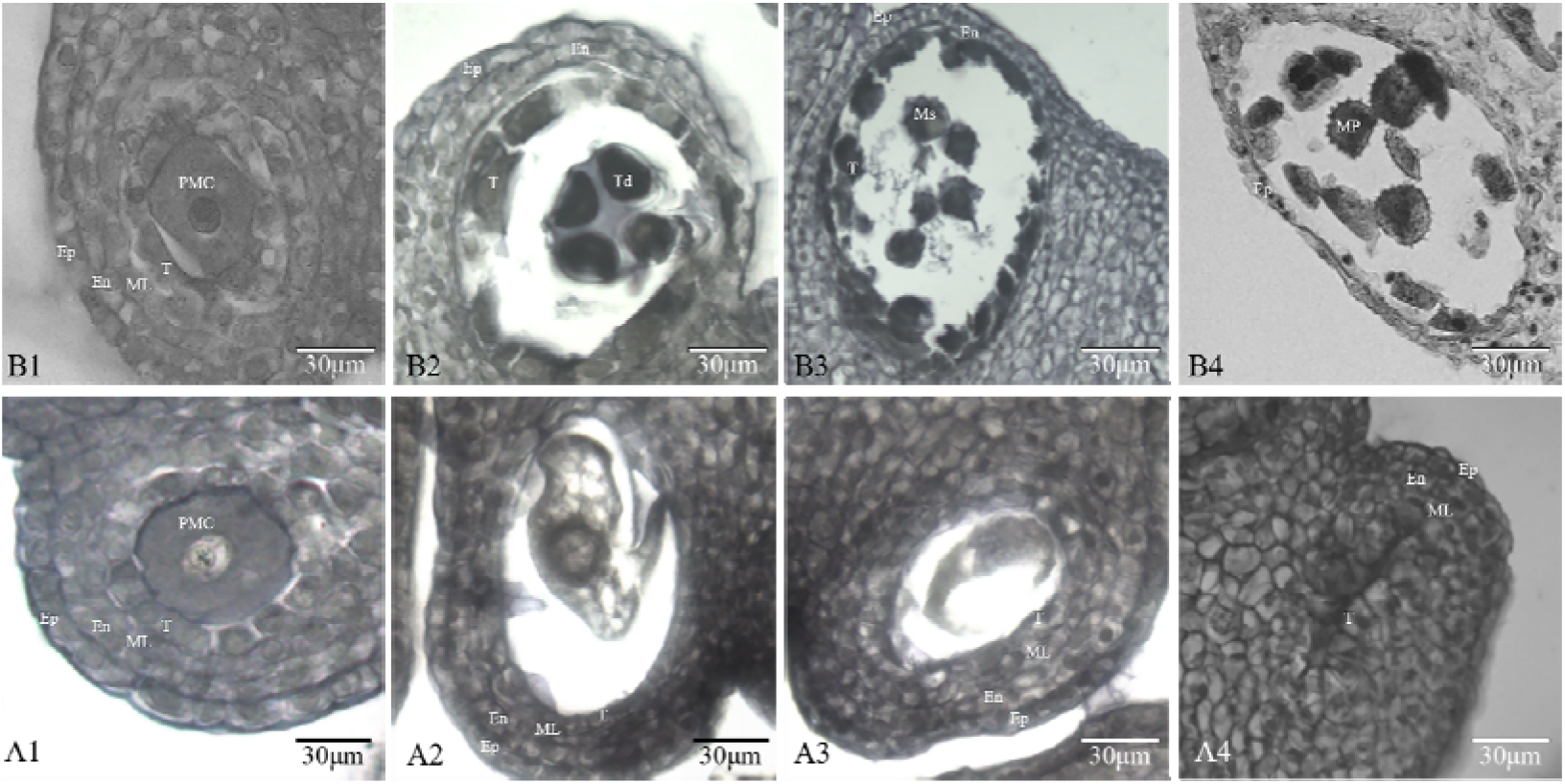
Comparative analysis of anther development between LD6A and LD6B. Notes: A1-A4, LD6A; B1-B4, LD6B; A1-B1, PMC; A2-B2, Td; A3-B3, early Uni; A4-B4, MP. PMC, pollen mother cell stage; Td, tetrad stage; early Uni, early uninucleate stage; MP, mature pollen stage. Ep, epidermis; En, endothecium; ML, middle layer; T, tapetum; Ms, microspore.

A microscopic analysis revealed no obvious differences in pollen size and morphology between CMS line and maintainer line at PMC stage (Figure1 A1, B1). Epidermis, endothecium, middle layer, tapetum and pollen mother cell are arranged closely at PMC stage. At tetrad stage (Figure1 A2, B2), the tetrad microspores can be observed clearly in LD6B, as middle layer was dissolved, and tapetum began to separate from anther wall and fragmentation. While in the CMS line, the four layer cells were closely connected without any obvious changes, but at the center of pollen sac, only highly vacuolated pollen mother cell was visible. With the pollen development, the tetrad microspores separate from each other and the tapetum cells were further condensed into a mass and stained deeper in maintainer line at the early uninucleate (Figure1 A3, B3, 5-6mm). Meanwhile, in the CMS line, only cell debris were remained in the pollen sac center. No visible changes were observed in the pollen wall of the two tested lines at this point. At MP stage (Figure1 A4, B4, > 6mm), pollen grains with spines were visible and only epidermis layer was present on the pollen wall in LD6B. By contrast, the pollen sac had become a solid linear structure in the LD6A, and the four pollen wall layers were still present as during the other stages.

Subcellular changes during pollen development were studied in detail with transmission electron microscopy (TEM) in CMS line and maintainer line. At PMC stage, the tapetum cells were closely linked with the other pollen wall layers and had a lighter staining in CMS line, while in maintainer line, tapetum cells began to separate from other layers and had a deeper stain (Figure3 A1 and B1). At tetrad stage, tapetum cells were degraded only in LD6B, but not in LD6A (Figure 3 A2 and B2). At MP stage, in LD6B, only epidermis and endothecium layers were present with spines of pollen grains were clearly visible (Figure 3A1). However, in LD6A, the four layers of pollen wall were completed, except cellular content were lost (Figure 3 B1).

**Figure 3.**
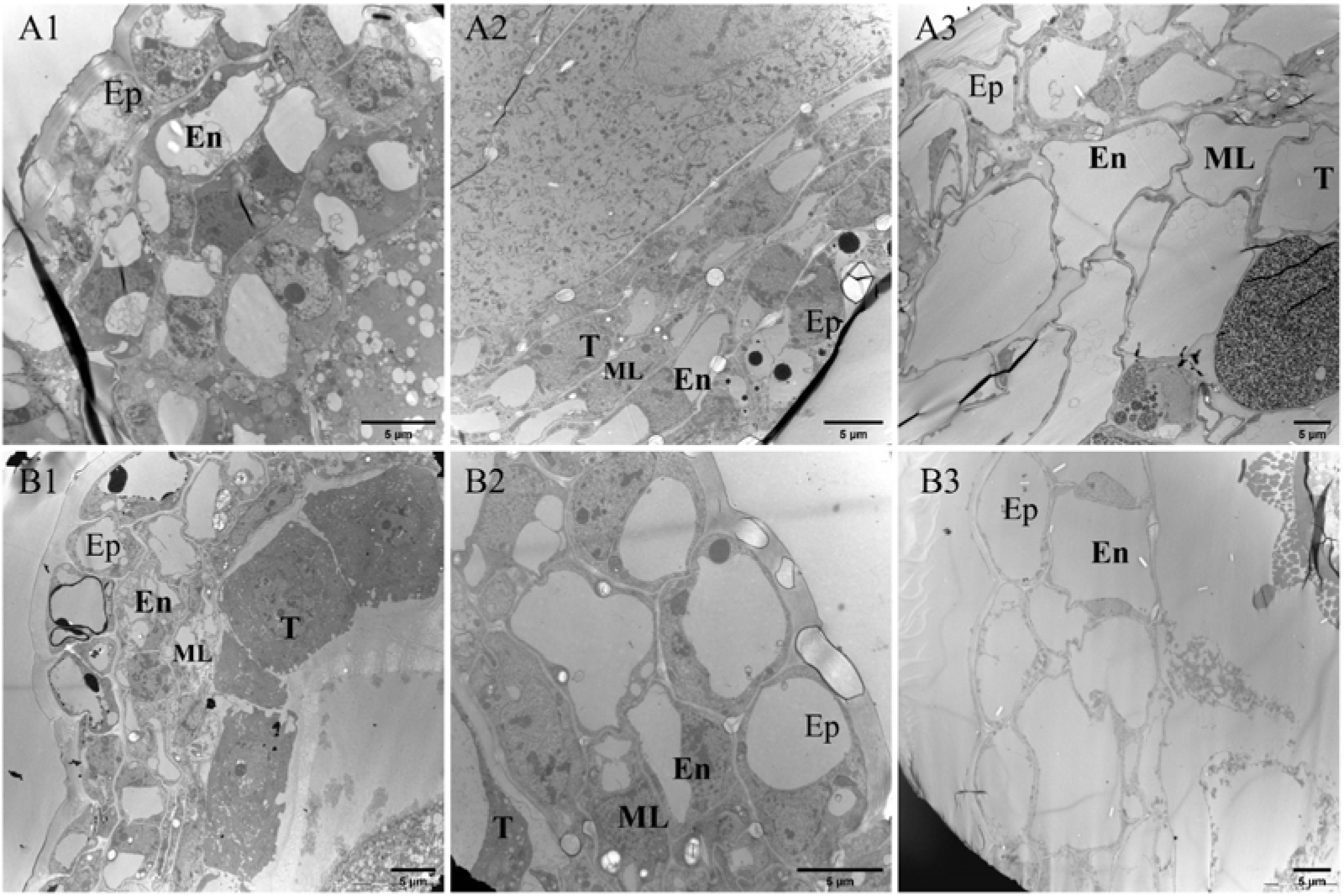
An overview of anther development by transmission electron microscope B1-B3, LD6B; A1-A3, LD6A; A1, B1, PMC; A2, B2, Td; A3, B3, MP. PMC, pollen mother cell stage; Td, tetrad stage; MP, mature pollen stage. Ep, epidermis; En, endothecium; ML, middle layer; T, tapetum.

Mitochondria are the energy supply sites for pollen development, and its structure had a great influence on pollen development. In LD6B, at PMC stage, mitochondrial cristae were not clearly visible due to stained tapetum, and mitochondria became invisible during MP stage. In CMS line, cristae were clearly observed before and after pollen abortion (PMC and MP stage) and stained lighter than Td stage (Figure 4). In the pollen abortion stage (Td stage), a great difference was observed between LD6A and LD6B. In LD6A, there were fuzzy and narrowly spaced cristae (Figure 4 A2), but cristae had high visibility in LD6B, besides, the space between cristae was similar with the normal conditions in LD6A (Figure 4, A1 and A3).

**Figure 4.**
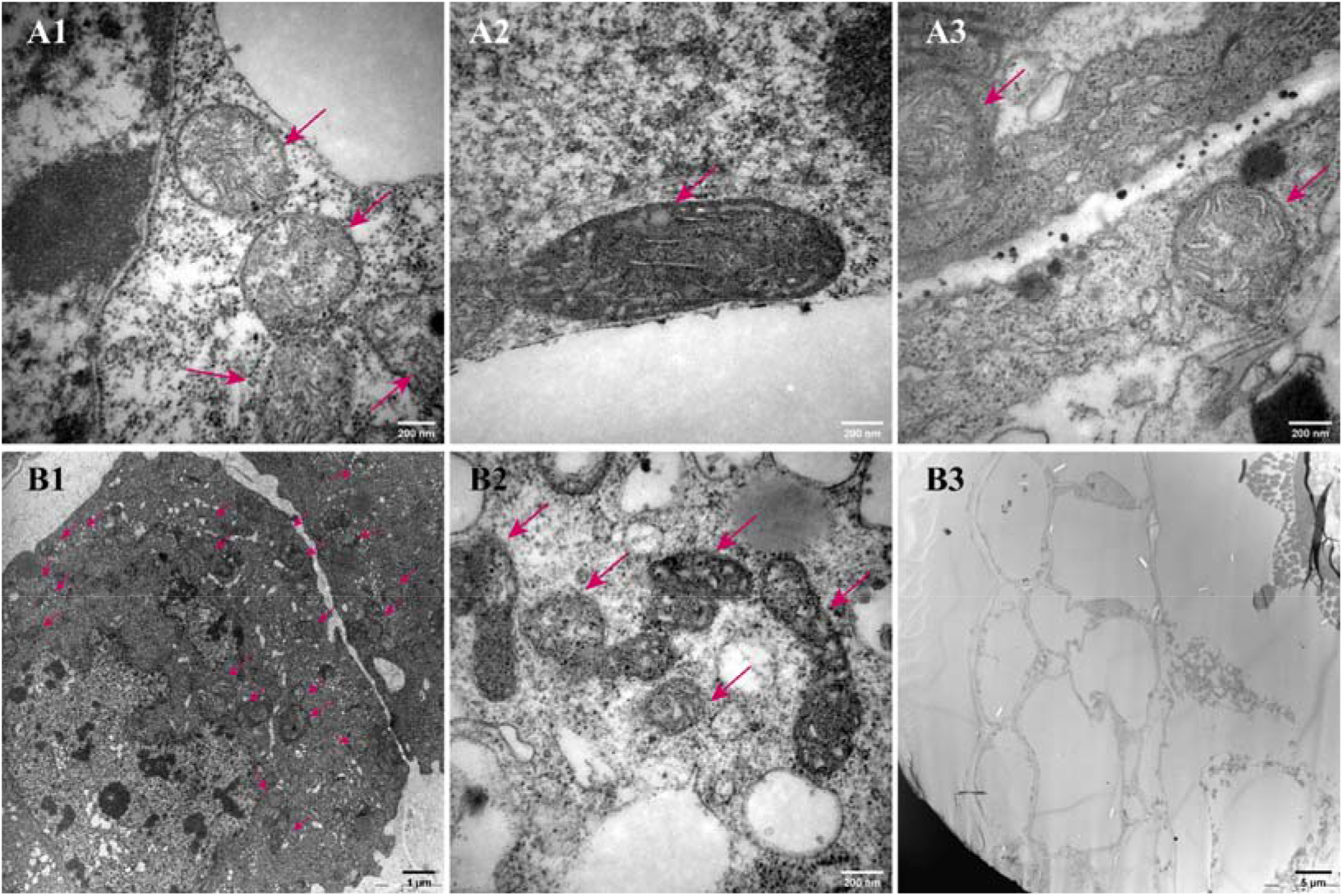
Comparative observation on mitochondrial ultrastructure in tapetum at pollen different developmental stages Notes: B1-B3, LD6B; A1-A3, LD6A; A1, B1, PMC; A2, B2, Td; A3, B3, MP. PMC, pollen mother cell stage; Td, tetrad stage; MP, mature pollen stage. Ep, epidermis; En, endothecium; ML, middle layer; T, tapetum. The arrow refers to mitochondria.

### 3.2 Antioxidant enzyme activity

Excessive ROS accumulation is an important signaling pathway in pollen development, which is reflected by an increased malondialdehyde (MDA) levels. LD6A contained significantly higher MDA content than LD6B at each pollen development stage (Figure 5d), indicating a significant damage to cellular membrane in CMS line.

**Figure 5.**
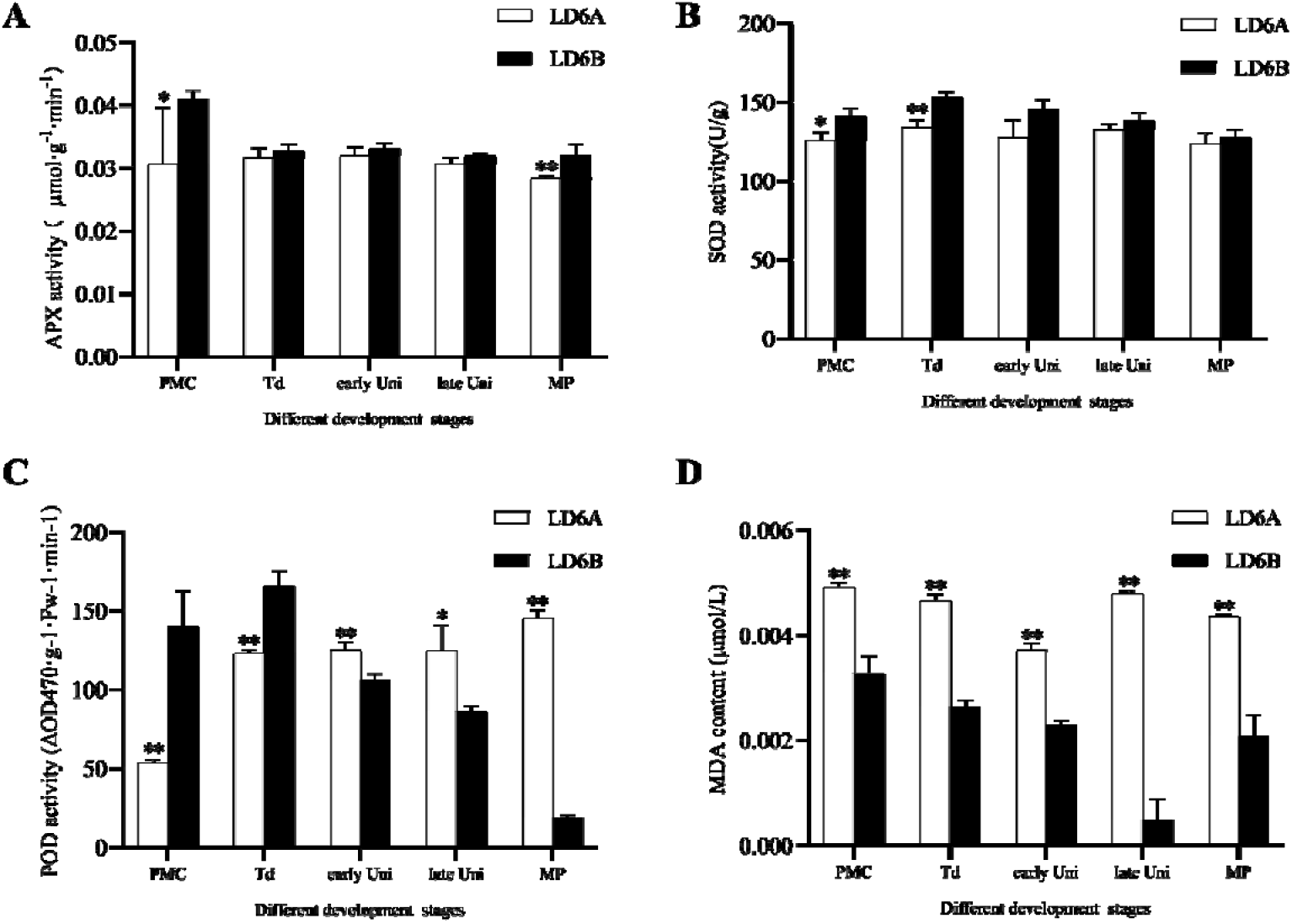
Antioxidant enzyme activity and MDA content of anthers at different developmental stages. A: APX activity; B: SOD activity; C: POD activity; D: MDA content. PMC, pollen mother cell stage; Td, tetrad stage; early Uni, early uninucleate; MP, mature pollen stage. * show significant difference between LD6A and LD6B at p□<□0.05 level, ** show remarkable significant difference at p□<□0.01 level, all the significance analyses were made by pairwise comparison in the same pollen development stage.

**Figure 6.**
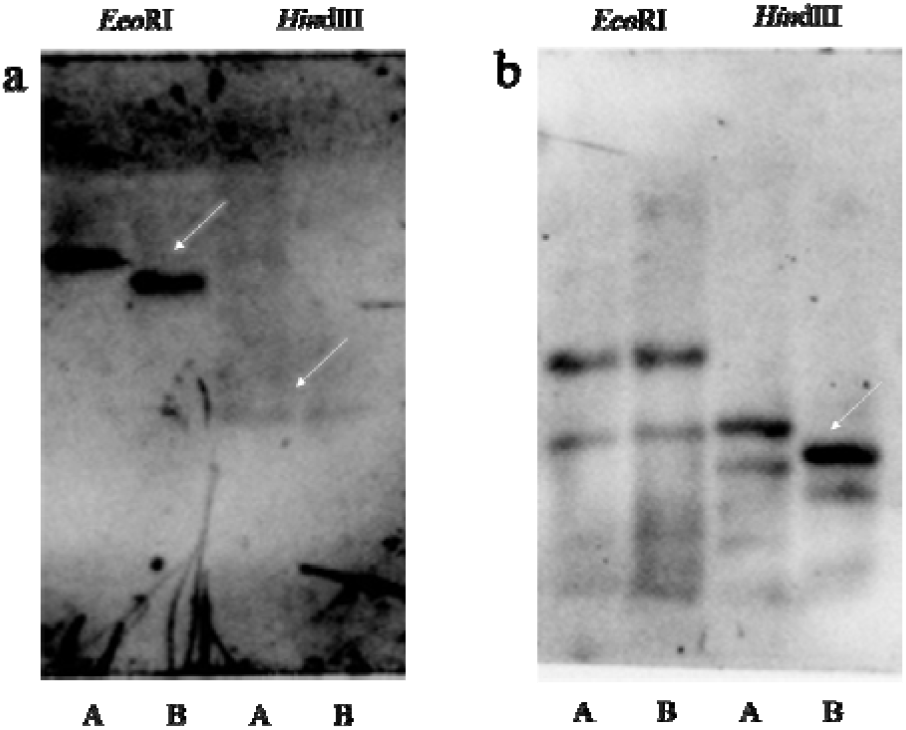
RFLP analysis of mitochondrial ribosomal protein genes *rps4* and *rpl10* Note: a, *rps4;* b, *rpl10;* A, CMS line LD6A; B, maintainer line LD6B

Activities of antioxidant enzymes such as APX, SOD, POD were significantly varied in CMS maintainer line during different developmental stages (i.e. PMC and Td stage, Figure 5 a, b and c). On average, CMS line has a relatively lower ability to capture unregulated ROS than its maintainer line.

For example, APX is a key enzyme to scavenge H_2_O_2_ in chloroplasts and cytoplasmic matrix(Wu *et al*., 2020), throughout pollen development, APX activity in CMS line was lower than its maintainer line, especially at PMC stage, when this difference was significant (Figure5A). Similarly, SOD (superoxide dismutase) plays an important role on the oxidation and antioxidant balance and is the first enzyme to regulate oxidative stress (Dias *et al*., 2019). SOD activity in CMS line became lower than maintainer line, particularly during PMC and Td stage (Figure 5B). POD (peroxidase) was considered as a physiological index of tissue aging (Nigam *et al*., 2019). CMS line had a significantly lower POD activity prior to microspore abortion (PMC and Td stage), but after pollen abortion, it was significantly induced. These opposite trends indicated a higher oxidative stress level before pollen abortion and a premature tissue aging after pollen abortion in CMS line.

### 3.3 RFLP analysis of mitoribosome protein genes

Mitoribosome is the core of protein translation in mitochondria, and it is made up of proteins and rRNAs (Waltz *et al*., 2020). RFLP analysis of mitochondrial ribosomal protein genes *rps4* and *rpl10* showed that *rps*4/*Eco*R□ had a large fragment in CMS line, while *rps4/Hin*d□ had very weak bands and same size in both CMS and maintainer lines. There was no polymorphism in *rpl*10/*Eco*R□, but two fragments of *rpl*10/*Hin*d□ were slightly larger in CMS than in its maintainer line. It showed mutations occurred at DNA level in or near the coding region of *rps*4 and *rpl10*, and impaired mitoribosome functioning in CMS line.

### 3.4 An overview of spectra and proteins functional annotation

Three random biological replicates and three technical duplicates experiment were detected from the floral buds of LD6A and LD6B in the proteomic analyses by iTRAQ technology. Totally 317,138 spectrums were generated, and 24,645 peptides and 7,024 proteins were identified with 1% FDR (Figure 7a) with a protein sequence coverage with 0-10%, 10-20%, 20-30%, 30-40%, 40-50%, 50-60%, >60% (Figure 7b).

**Figure 7.**
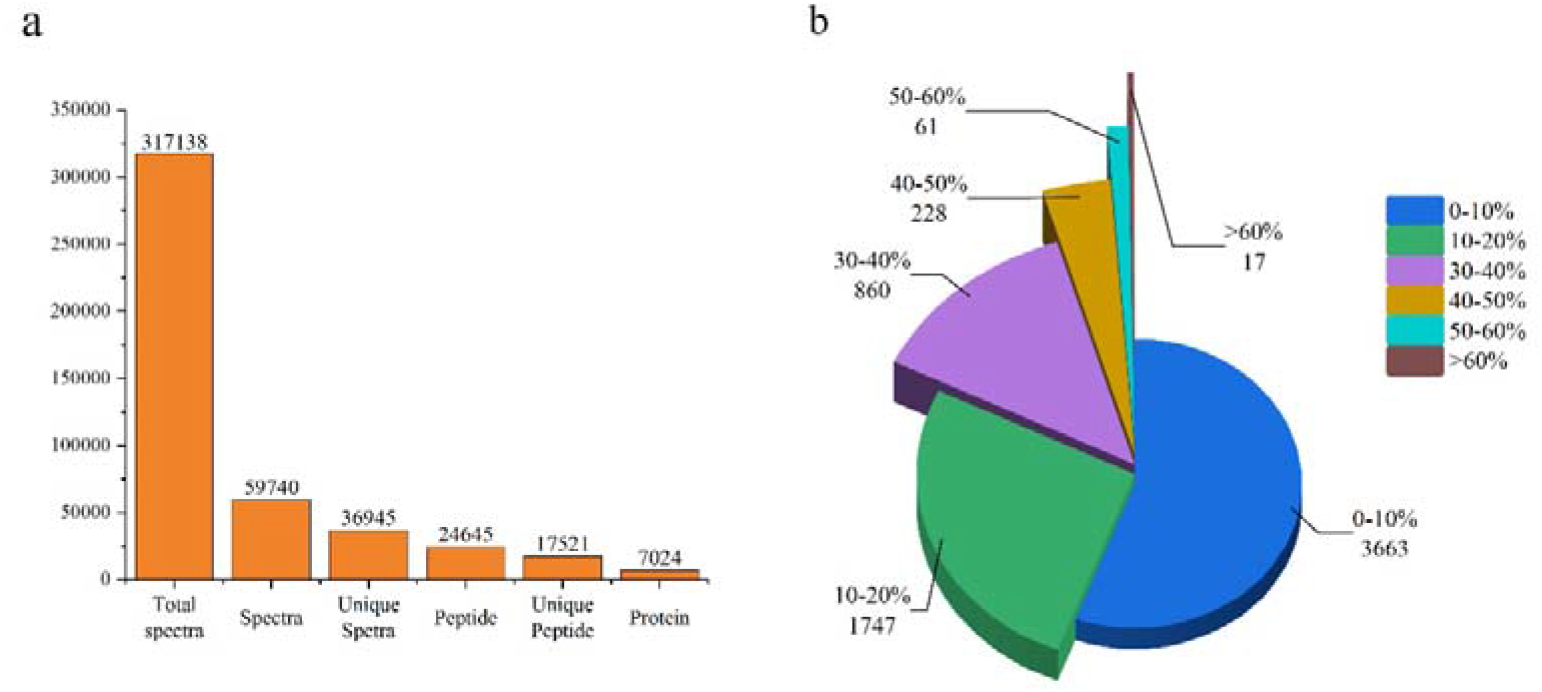
Overview of protein identification and distribution of protein coverage Note: a, overview of protein identification; b, distribution of protein coverage

**Figure 8.**
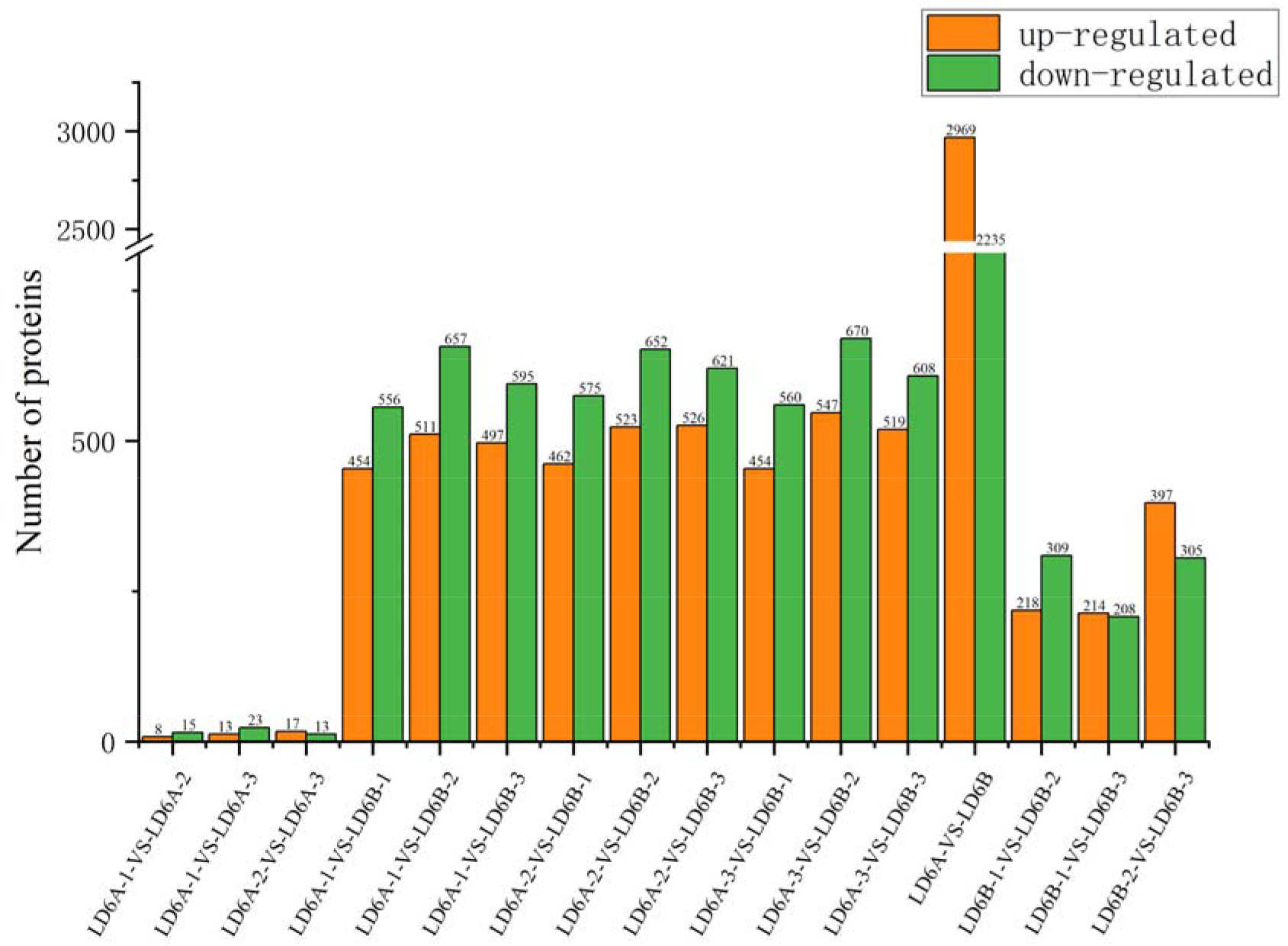
Number of differentially expressed proteins in different groups

### 3.5 Proteins quantification and annotation

In this project, we set three repeats of LD6A and LD6B as comparison groups. The final differentially expressed proteins repeat experiment were defined with 1.2-fold change (mean value of all comparison groups) and P-value (t-test) less than 0.05. In A to A and B to B comparison groups, the numbers of DEPs are very close. In each A to B comparison groups, up-regulated and down-regulated figures are basically consistent. In totally comparison group, 2969 up-regulated and 2235 down-regulated proteins are identified.

Gene Ontology, or GO, is a major bioinformatics initiative to unify the representation of gene and gene product attributes across all species. Membrane related GO terms in cellular component, organism development associate terms in biological process, as well as electron carrier activity and antioxidant activity in molecular function predicated vigorous vital activities in membrane construction, pollen development and ROS balance (Figure 9).

**Figure 9.**
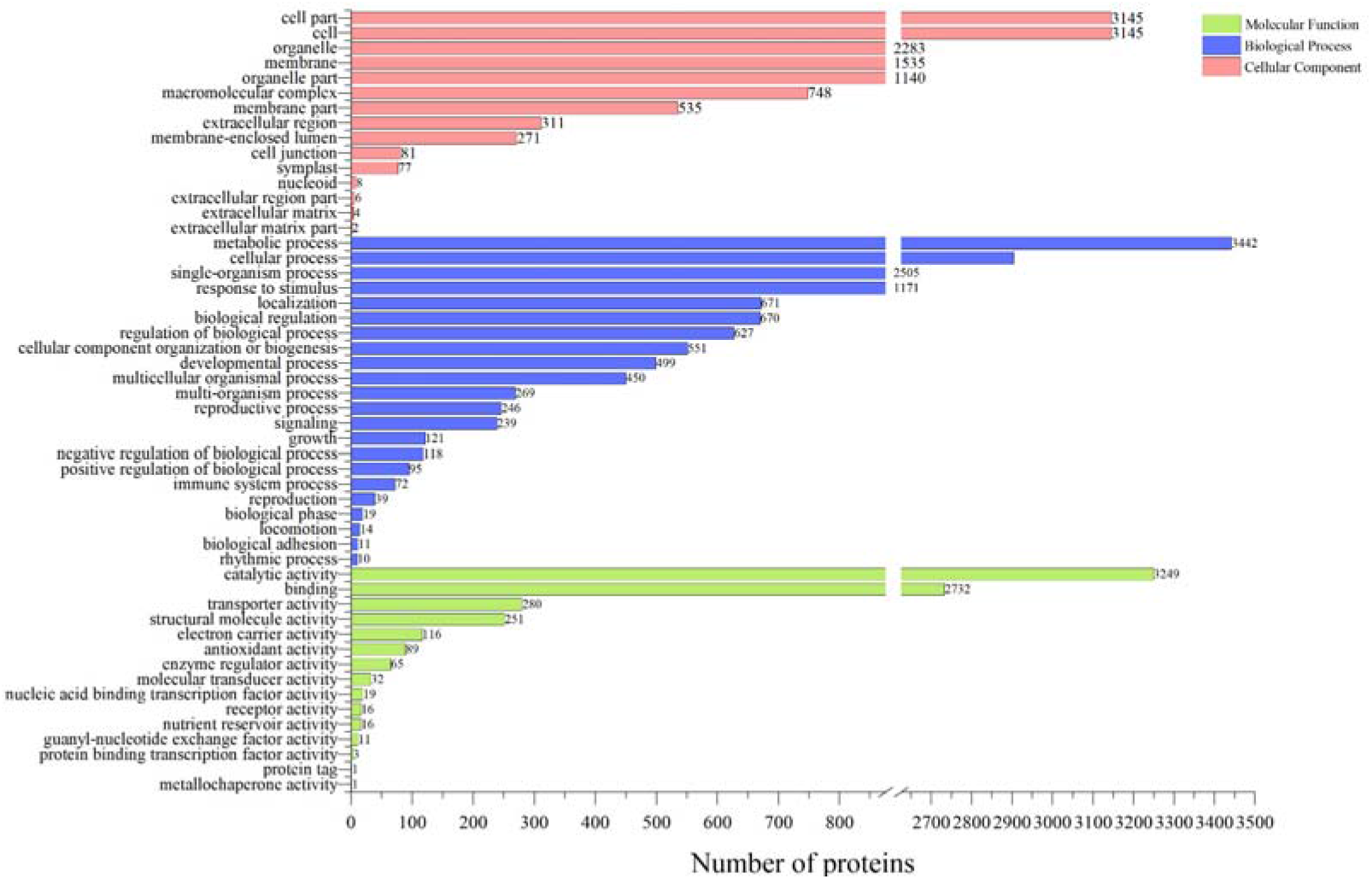
Gene ontology annotation analysis

Clusters of Orthologous Groups of proteins (COGs) were delineated by comparing protein sequences encoded in complete genomes, representing major phylogenetic lineages. Ribosome structure, protein turnover, cell wall/membrane/envelope biogenesis takes up a p high proportion (Figure 10).

**Figure 10.**
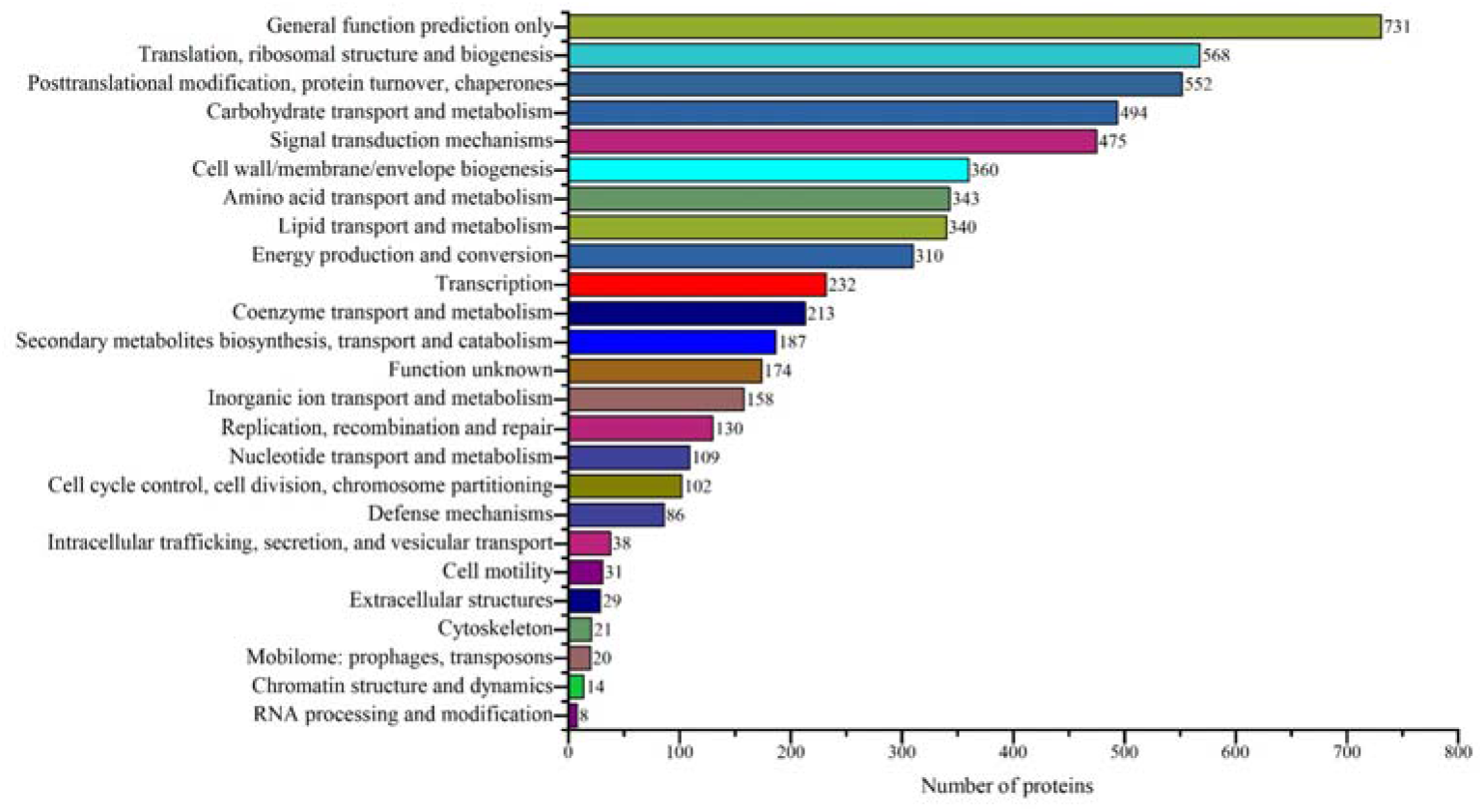
COG annotation analysis.

### 3.6 Differently Expression Proteins Analysis

Proteins usually interact with each other to regulate certain biological functions. We performed a pathway enrichment analysis of the differentially expressed proteins using KEGG database for each pairwise comparison (Figure 11). KEGG pathways for starch and sucrose metabolism, ribosome, proteasome, oxidative phosphorylation, and nucleotide excision repair were significantly difference (P<0.05) of the pathway enrichment in DEPs (Figure 11). In addition, we list the top 25 enriched KEGG pathways as Table 1.

**Figure11.**
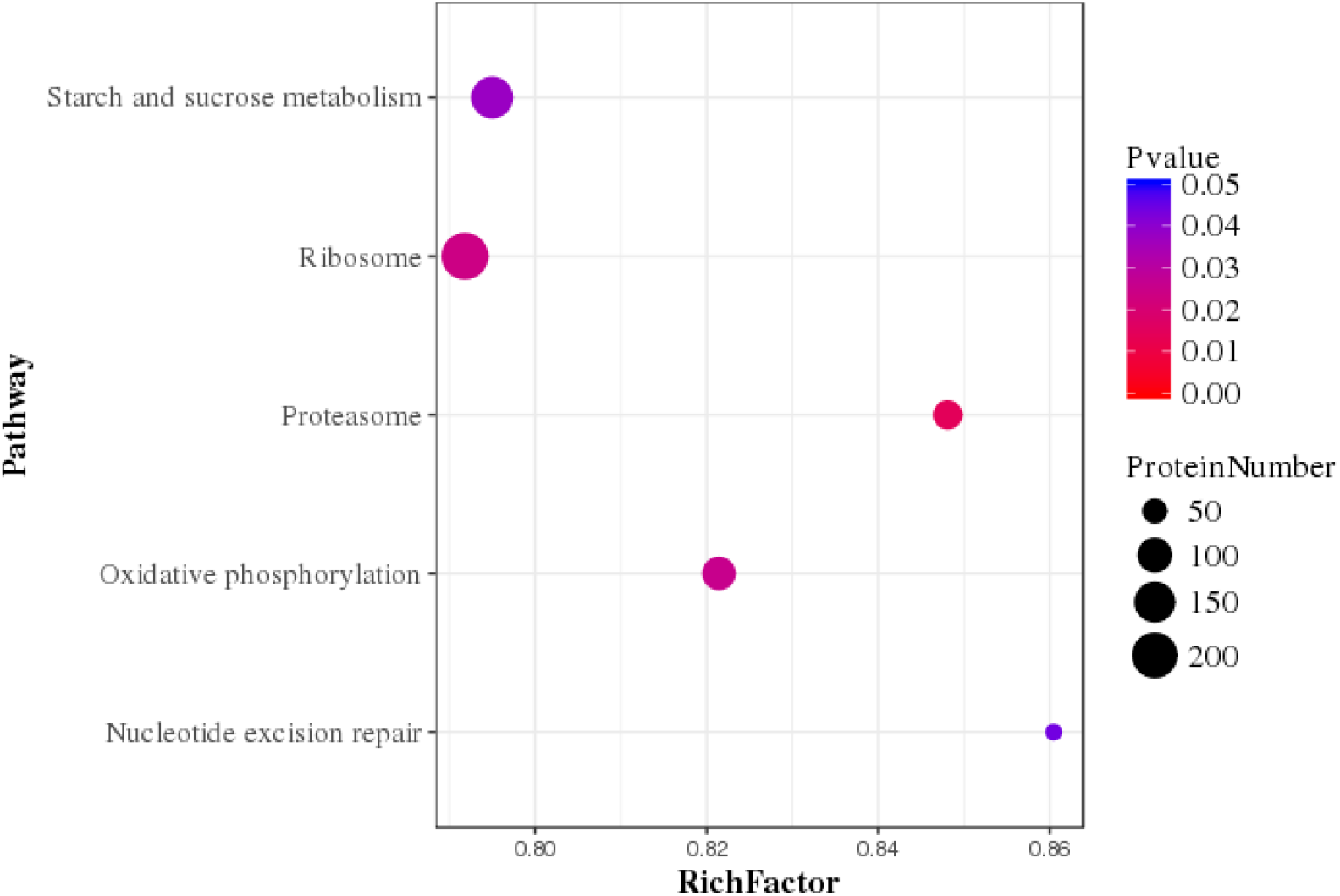
Statistics of pathway enrichment of differentially expressed proteins in LD6A vs LD6B pairwise.

**Table 1.**
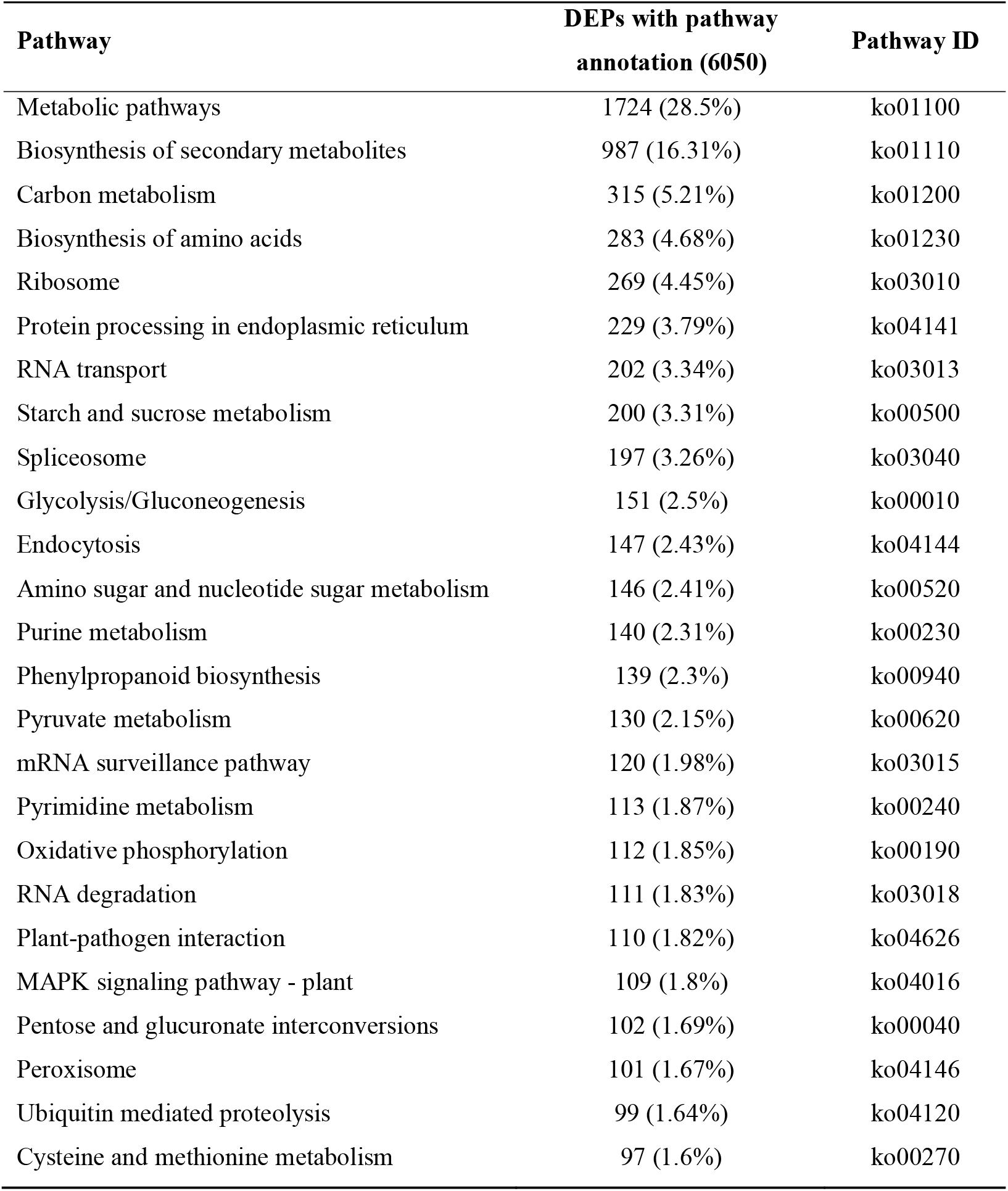
Statistical DEPs annotation analysis of Kyoto Encyclopedia of Genes and Genomes (KEGG) metabolic pathways

| Pathway | DEPs with pathway<br>annotation (6050) | Pathway ID |
| --- | --- | --- |
| Metabolic pathways | 1724 (28.5%) | ko01100 |
| Biosynthesis of secondary metabolites | 987 (16.31%) | ko01110 |
| Carbon metabolism | 315 (5.21%) | ko01200 |
| Biosynthesis of amino acids | 283 (4.68%) | ko01230 |
| Ribosome | 269 (4.45%) | ko03010 |
| Protein processing in endoplasmic reticulum | 229 (3.79%) | ko04141 |
| RNA transport | 202 (3.34%) | ko03013 |
| Starch and sucrose metabolism | 200 (3.31%) | ko00500 |
| Spliceosome | 197 (3.26%) | ko03040 |
| Glycolysis/Gluconeogenesis | 151 (2.5%) | ko00010 |
| Endocytosis | 147 (2.43%) | ko04144 |
| Amino sugar and nucleotide sugar metabolism | 146 (2.41%) | ko00520 |
| Purine metabolism | 140 (2.31%) | ko00230 |
| Phenylpropanoid biosynthesis | 139 (2.3%) | ko00940 |
| Pyruvate metabolism | 130 (2.15%) | ko00620 |
| mRNA surveillance pathway | 120 (1.98%) | ko03015 |
| Pyrimidine metabolism | 113 (1.87%) | ko00240 |
| Oxidative phosphorylation | 112 (1.85%) | ko00190 |
| RNA degradation | 111 (1.83%) | ko03018 |
| Plant-pathogen interaction | 110 (1.82%) | ko04626 |
| MAPK signaling pathway - plant | 109 (1.8%) | ko04016 |
| Pentose and glucuronate interconversions | 102 (1.69%) | ko00040 |
| Peroxisome | 101 (1.67%) | ko04146 |
| Ubiquitin mediated proteolysis | 99 (1.64%) | ko04120 |
| Cysteine and methionine metabolism | 97 (1.6%) | ko00270 |

### 3.7 Integrated candidate CMS-related proteins, ROS reduce enzymes protein expression profile analysis

In earlier studies, many CMS associated candidate genes were screen out through transcriptome analysis, such as *Abrin, HAT, MDH, atp8* (Zheng *et al*., 2019). After annotation and profile analysis, the upregulation and downregulation trend were same between transcriptome and proteome (Figure 12). At the same time, we annotated some enzymes related to TCA, such as IDH and ME, and found results showed a downregulation trend as well as MDH. Some ROS reduction isoenzymes, like SOD, POD, APX, showed a general downregulation trend (Figure 13).

**Figure 12.**
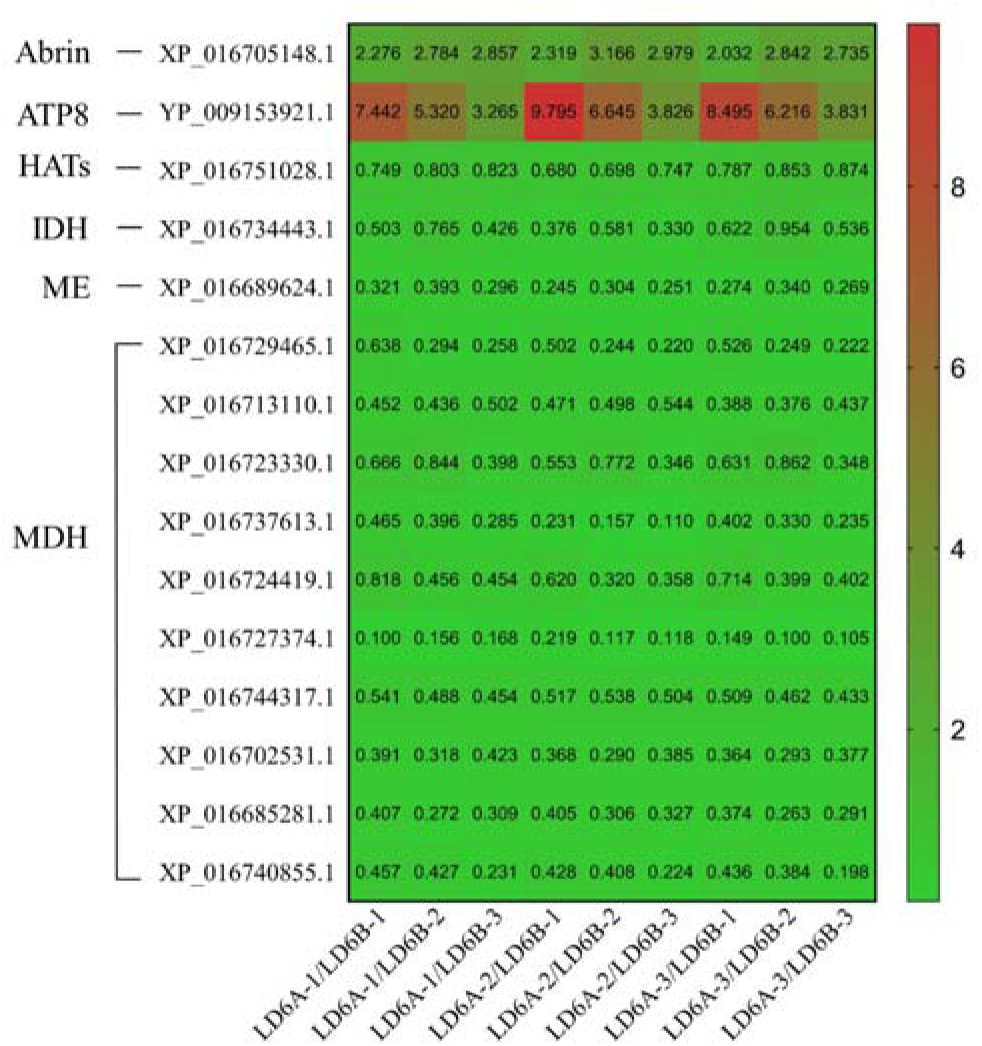
Relative expression of proteins associated CMS (Fold Change)

**Figure 13.**
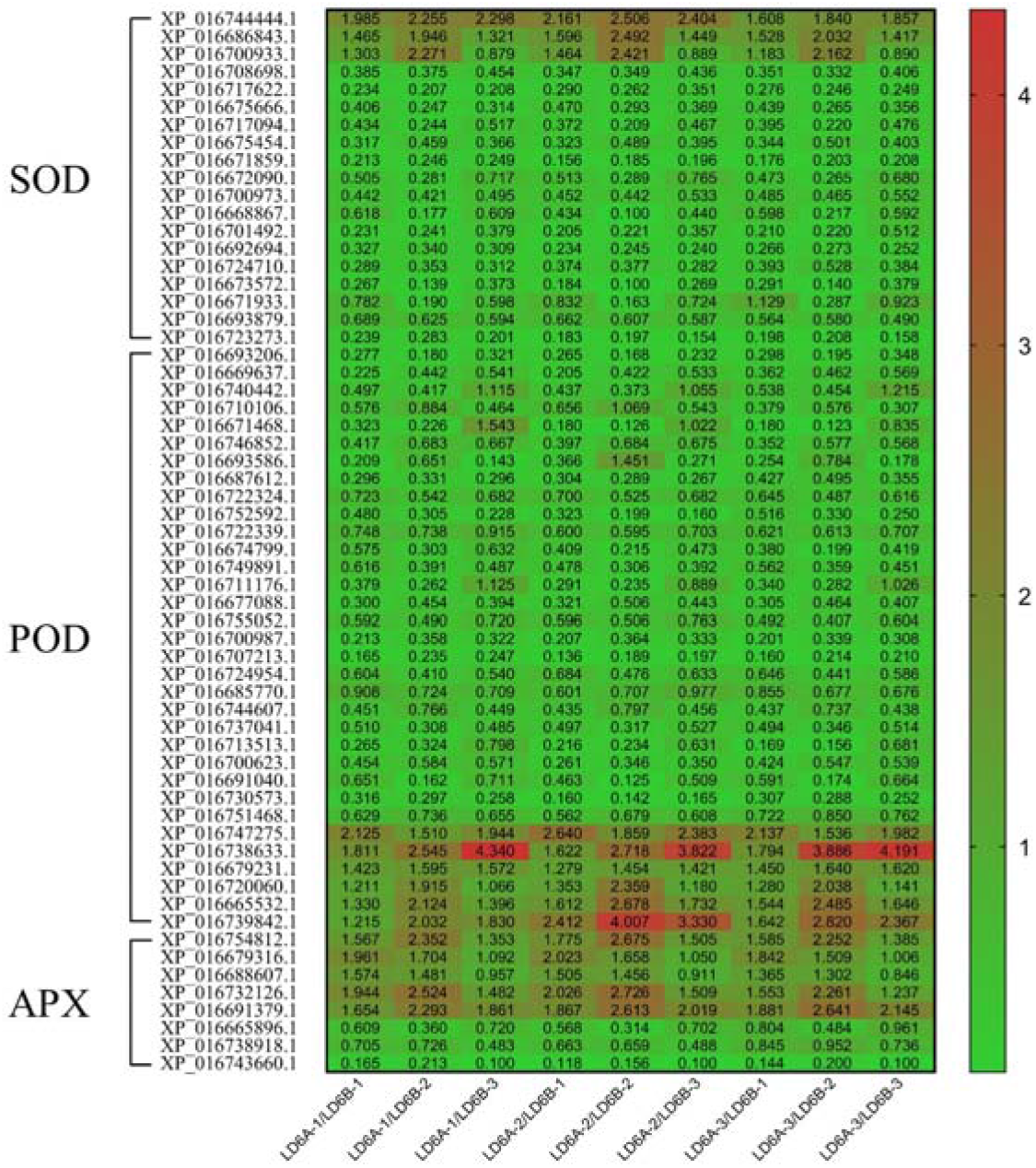
Relative expression of enzymes related to reactive oxygen species metabolism (Fold Change)

### 3.8 Validation of DEPs by qRT-PCR

qRT-PCR was used to verify the reliability of iTRAQ in RNA level, 12 DEPs (6 upregulated and 6 downregulated) were selected randomly. The results confirmed the reliability of iTRAQ data (Table 2). The fold change of some DEPs confirmed by qRT-PCR was different to that was detected by iTRAQ, such as XP_016722577.1, XP_016678431.1, XP_016750824.1, XP_016746825.1, even regulated trend is opposite. This could be due to different expression model between RNA and protein. To explore further expression model of interesting proteins, qRT-PCR at different pollen development stage were detected. As Figure14 shown, the expression model of every protein was different.

**Table 2.** qRT-PCR confirmation of the expression profiles of selected proteins

| Protein ID | Description | Fold Change |  |
| --- | --- | --- | --- |
|  |  | iTraq | qRT-PCR |
| XP_016735888.1 | ADP, ATP carrier protein 3, mitochondrial | 8.74 | 6.69 |
| XP_016722577.1 | histone H4-like | 7.69 | 0.44 |
| XP_016729156.1 | V-type proton ATPase subunit F isoform X4 | 7.30 | 5.58 |
| XP_016724302.1 | cytochrome P450 CYP749A22-like | 7.20 | 4.38 |
| XP_016678431.1 | ATPase 10, plasma membrane-type-like | 4.72 | 0.65 |
| XP_016694288.1 | cytochrome c oxidase subunit 5C-like | 5.55 | 1.01 |
| XP_016676713.1 | anther-specific protein LAT52-like | 0.11 | 0.79 |
| XP_016684521.1 | pollen allergen Che a 1-like | 0.14 | 0.13 |
| XP_016750824.1 | ADP, ATP carrier protein 1, mitochondrial-like | 0.21 | 49.0 |
| XP_016746825.1 | mitochondrial outer membrane protein porin of 34 kDa-like | 0.22 | 2.18 |
| XP_016745836.1 | triacylglycerol lipase 2-like isoform X2 | 0.16 | 0.28 |
| XP_016750648.1 | acyl-coenzyme A thioesterase 13-like | 0.18 | 0.41 |

**Figure 14.**
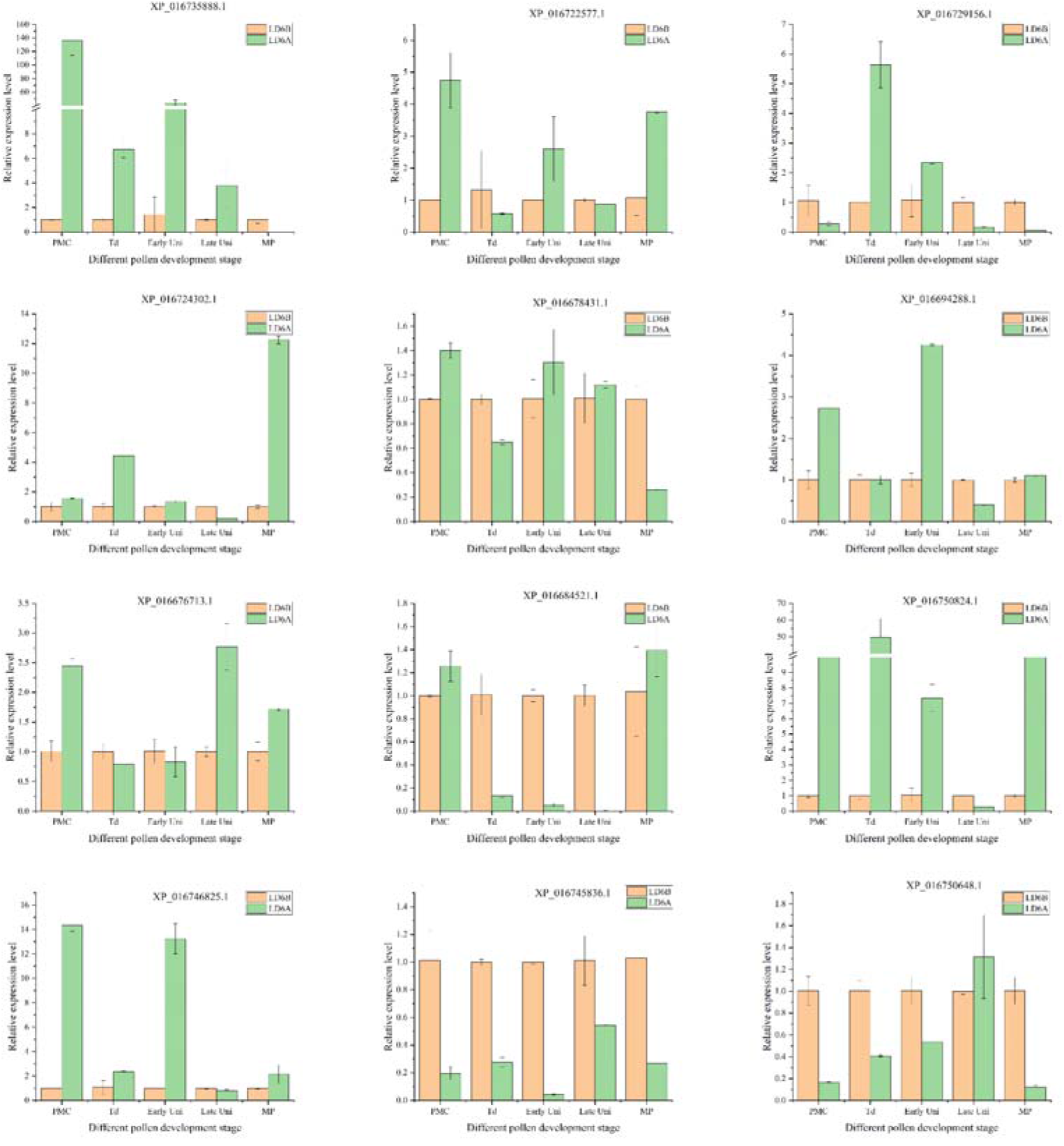
Expression profiles of selected proteins at different pollen development stage

**Figure 15.**
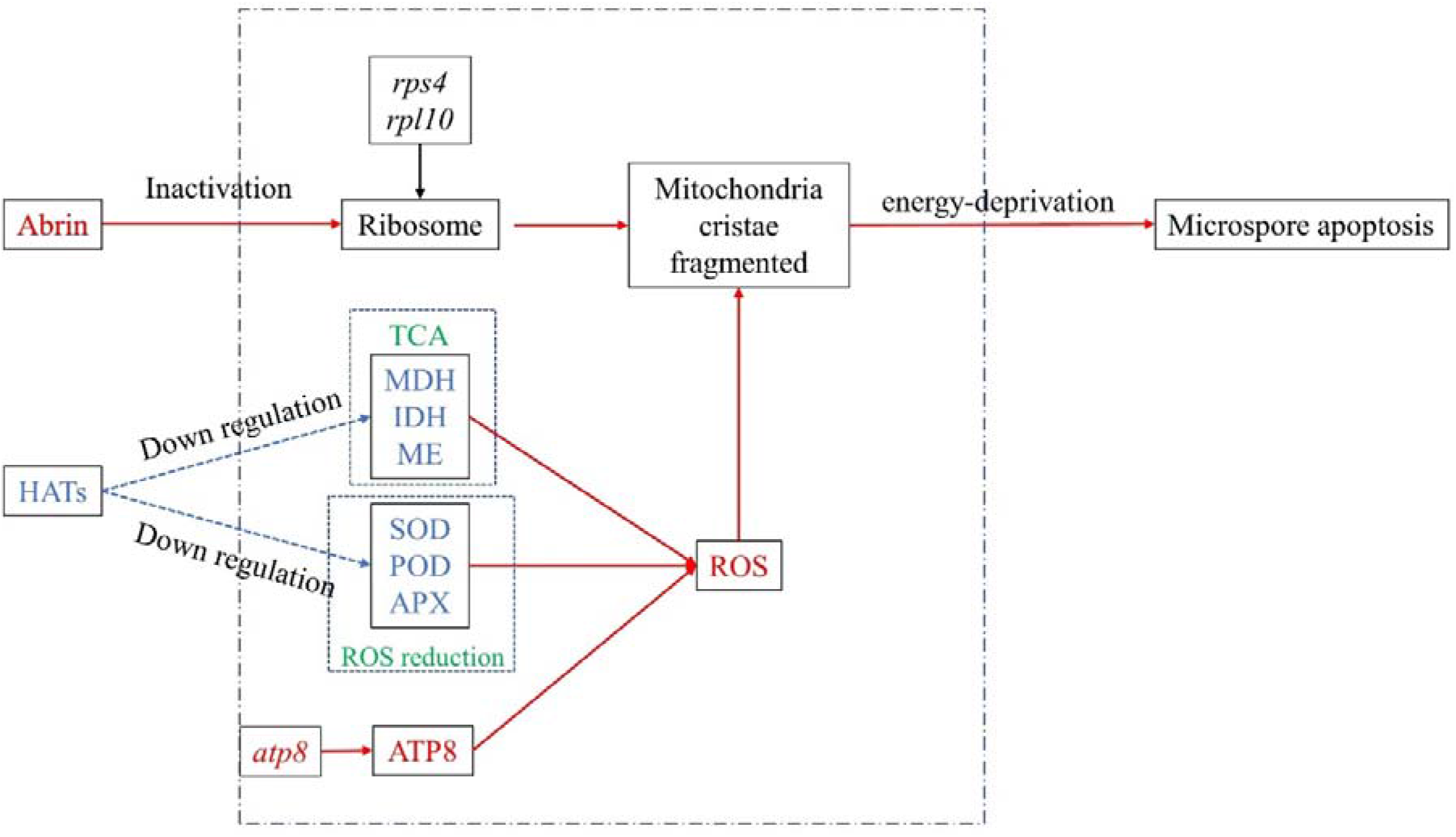
Schematic diagram of LD6A microspore abortion hypothesis (Red indicates promotion or up regulation; blue indicates inhibition or down regulation)

## 4 Discussion

### 4.1 The non-degradation of tapetum resulted in the abortion of microspore

Mitochondria are the main cellular sites for energy conversion in plants tissues (Rhiannon R. White, 2020). Energy conversion process include ATP synthesis on mitochondrial complex V after glycolysis, tricarboxylic acid cycle (TCA) and oxidative phosphorylation (Etienne Meyer, 2019). Without timely and sufficient energy supply, many normal developmental processes including pollen development are impaired, leading to cytoplasmic male sterility. In cotton anther, four layers of pollen sac cells are arranged around microspores, whereas tapetum cells, which provide energy for microscope development through degradation. Our study indicated that in LD6A, tapetum layer cells remained intact during each pollen development stage, however microspore disappeared after Td stage (Fig2 and Fig3). At the same time, in LD6B, tapetum layer cells degraded gradually, as microspore develop into mature pollen. This indicates that poor energy supply from the non-degraded tapetum led to microspore abortion in LD6A.

### 4.2 The abnormality of mitochondrial ribosomal genes may affect the structural mitochondria

Mitochondria contain a separate genome with fully functional gene expression machinery, even though most proteins acting in mitochondria (>95%) are nuclear encoded (Giegé *et al*., 2005). Mitochondrial ribosomes (mitoribosome) strongly influence the inner membrane structure through translation mechanism (Pfeffer *et al*., 2015b). This inner membrane is known as the location of electron transfer chain (ETC), and base of mitochondrial functioning. Genes such as *rps4* and *rpl10* encode two subunits of mitoribosome. RFLP polymorphisms between LD6A and LD6B (Fig 6) revealed possible mutations at DNA level in or near the coding region of *rps4* and *rpl10*, which altered translation level of mitroribosome. At protein level, a significant DEPs number in ribosome pathway (Figure 12) indicates possible DNA level mutations in *rps4* and *rpl10* influenced the protein expression level in mitroribosome. Furthermore, these potential changes affected mitochondrial structure, as evidenced by TEM (Fig4b, e).

### 4.3 The burst of ROS affected the energy supply of microspore development

Oxidative stress and abnormal cellular ROS can deplete ATP and NADH by blocking ascorbic acid glutathione cycle of mitochondria (Ling Huang, 2011) and degrading mitochondrial genomic DNA (Wan *et al*., 2007). ROS can also cause plasma membrane peroxidation, thus affecting the function and structure of the membrane (Abdelrahman *et al*., 2020; Dias & Nylandsted, 2021). In this study, upregulated genes *Abrin*, *atp8* may induce ROS generation in the CMS line, while ROS regulation was weakened due to downregulation of MDH - a key TCA enzyme gene. Similarly, a histone acetyltransferase gene HAT can regulate gene expression in chrome level (Wang *et al*., 2014), which is upregulated in CMS line. Based on the comparative analysis of the related indexes of active oxygen metabolism between CMS and maintainer line, we can speculate that from pollen mother cell, activity of key antioxidant enzymes (e.g. APX, SOD and POD) in anther of CMS line were significantly decreased (Fig. 4-1), and thus ability to eliminate ROS. This was further supported by a significant increase in MDA (Fig. 4-1, d), suggesting structural damage to mitochondrial membrane in CMS line. Subcellular microstructure observations of anther mitochondria of CMS line also indicated fuzzy ridge and narrow plasma membrane space (Fig. 2-2). This impaired structural and functional of mitochondria curtailed normal energy conversion and depletion of ATP and NADH. Poor mitochondrial capacity of converting energy produced from glycolysis into ATP, could induce ROS accumulation, a by-product of glycolysis. This was exacerbated by lack of energy supply from non-degrading tapetum cells which impaired micropore development (Fig. 2-4).

### 4.4 Aberrant high expression of Abrin results in microspore apoptosis

In this study, Abrin, a toxic protein, have a higher expression in RNA and protein level at Td stage in CMS line (Fig13) which can induce mitochondrial apoptosis through ribosomal pathway in cancer cells (Narayanan *et al*., 2004). The production of toxic proteins in anthers could also induce CMS (Levings, 1993). The protein interaction database (DIP, https://dip.doe-mbi.ucla.edu/dip/Main.cgi) shows two proteins interact with Abrin. A glycosylation related protein, udp-n-acetyl-d-galactosamine: polypeptide n-acetylgalactosyltransferase (gly5, DIP: 26207n), which is located on the Golgi body; The other is acetylglutamate kinase (NAGK, DIP: 5348n), which is a catalytic enzyme in the second step of arginine biosynthesis in plastids and regulates gametophyte function and embryonic development (Huang *et al*., 2017).

### 4.5 A hypothesis of CMS molecular mechanism

Integrated comparative analysis of cytology, mitochondrial genome, reactive oxygen species metabolism, transcriptome and proteome, we found some interrelated results. For example, subcellular changes such as poor degradation of tapetum layer cells and mitochondrial membrane damage in CMS indicate energy starvation for developing pollen. At gene level RFLP polymorphism in mitochondrial ribosome genes *rps4* and *rpl10*, and Abrin could lead to ribosome inactivation in CMS line. At functional level, downregulation of antioxidant enzymes and subsequent MDA accumulation suppressed TCA related enzymes such as MDH, IDH and ME. Based on these results, we propose the following molecular mechanism hypothesis of CMS in cotton: at chromatin level, the inhibition of *HAT* (histone acetyltransferase) expression down-regulated series genes. Upregulation of *atp8* caused a large consumption of H^+^ in the electron transport chain and a leak of electronics, which increased the accumulation of ROS. These factors lead to ROS outbreak in tapetum cell. Furthermore, ROS destroyed proteins structure on the mitochondrial membrane. On the other hand, the up-regulated Abrin inactivated the ribosome in mitochondrion. But the muted of *rps4* and *rpl10* (mitochondrial ribosome protein) cannot supplement the functional mitochondrial ribosome proteins. The destruction of structural proteins by ROS and the disconnection of structural proteins by ribosome inactivation led to the abnormality of mitochondrial structure. The disabled mitochondria could not transform the energy of tapetum cells, so that microspores could not get enough energy supply and abortion.

## 5 Conclusions

In this study, cytological, mitochondrial genome, ROS metabolism, transcriptomic and proteomic analyses were compared between the CMS line LD6A and its maintainer line LD6B. It was found that LD6A microspore was aborted at Td stage but tapetal cells did not degrade during anther development as pollen sacs collapsed into a line. Further, the inner ridge of mitochondria in tapetum cells of LD6A at Td stage was fuzzy and membranous space was narrow. Antioxidant enzymes such as APX and SOD in CMS line had lower activities than those in maintainer lines during anther development; POD activity of male sterile line was lower than that of maintainer line before anther abortion, but higher than that of maintainer line after anther abortion; Due to poor regulation of antioxidant enzymes, CMS line could not eliminate ROS and and thus suffered membrane damage. The expression correlation of Abrin, HAT, ATP8 and MDH was analyzed by multi omics and multi-dimensional comparative analysis. The results of transcriptome and proteome were verified by each other. The physiological indexes and proteomic results had verified each other. Our findings will give a new insight into cotton CMS mechanism, which could improve utilization of heterosis by CMS line in cotton breeding program.

## Author Contributions

J.Z. performed the experiments and drafted the manuscript. A.K., B.Z., X.K. participated in the experiments. A.K., Z.Q., U.N., Y.L. participated in the revision of the manuscript. F.L., R.Z. conceived, designed, and supervised the study. All authors read and approved the final manuscript.

## Acknowledgements

We acknowledge Ming Tinghui (graduated from our team as a master) for her earlier basic morphology research on LD6A.

## References

Abdelrahman M, Ishii T, El-Sayed M, Tran LP. 2020. Heat sensing and lipid reprograming as a signaling switch for heat stress responses in wheat. PLANT AND CELL PHYSIOLOGY 61(8): 1399–1407.

Daniele, Del, Rio, And, Amanda, J., Stewart, And, Nicoletta, Pellegrini. 2005. A review of recent studies on malondialdehyde as toxic molecule and biological marker of oxidative stress. Nutrition Metabolism & Cardiovascular Diseases.

Dewey RE, Timothy DH, Levings CS. 1987. A mitochondrial protein associated with cytoplasmic male sterility in the t cytoplasm of maize. PROCEEDINGS OF THE NATIONAL ACADEMY OF SCIENCES OF THE UNITED STATES OF AMERICA 84(15): 5374–5378.

Dias C, Nylandsted J. 2021. Plasma membrane integrity in health and disease: Significance and therapeutic potential. Cell Discovery 7(1): 4.

Dias MC, Mariz-Ponte N, Santos C. 2019. Lead induces oxidative stress in Pisum sativum plants and changes the levels of phytohormones with antioxidant role. PLANT PHYSIOLOGY AND BIOCHEMISTRY 137: 121–129.

Etienne H. Meyer EWAC. 2019. Assembly of the complexes of the oxidative phosphorylation system in land plant mitochondria. Annual Review of Plant Biology.

Giegé P, Sweetlove LJ, Cognat V, Leaver CJ. 2005. Coordination of nuclear and mitochondrial genome expression during mitochondrial biogenesis in arabidopsis. The Plant Cell 17(5): 1497–1512.

Gupta PK, Balyan HS, Gahlaut V, Saripalli G, Pal B, Basnet BR, Joshi AK. 2019. Hybrid wheat: Past, present and future. THEORETICAL AND APPLIED GENETICS 132(9): 2463–2483.

H H, JM G, JM G. 1995. Characterization of the mitochondrial orf B gene and its derivative, orf 224, a chimeric open reading frame specific to one mitochondrial genome of the “ Polima “ male-sterile cytoplasm in rape seed (Brassica napus L.). CURRENT GENETICS 6(28): 546–552.

Hu J, Huang W, Huang Q, Qin X, Yu C, Wang L, Li S, Zhu R, Zhu Y. 2014. Mitochondria and cytoplasmic male sterility in plants. MITOCHONDRION 19: 282–288.

Huang J, Chen D, Yan H, Xie F, Yu Y, Zhang L, Sun M, Peng X. 2017. Acetylglutamate kinase is required for both gametophyte function and embryo development inArabidopsis thaliana. Journal of Integrative Plant Biology 59(9): 642–656.

Jing B, Heng S, Tong D, Wan Z, Fu T, Tu J, Ma C, Yi B, Wen J, Shen J. 2012. A male sterility-associated cytotoxic protein ORF288 in Brassica juncea causes aborted pollen development. JOURNAL OF EXPERIMENTAL BOTANY 63(3): 1285–1295.

Kong X, Liu D, Liao X, Zheng J, Diao Y, Liu Y, Zhou R. 2017a. Comparative analysis of the cytology and transcriptomes of the cytoplasmic male sterility line H276A and its maintainer line H276B of cotton (Gossypium barbadense l.). INTERNATIONAL JOURNAL OF MOLECULAR SCIENCES 18(11): 2240.

Kong X, Liu D, Liao X, Zheng J, Diao Y, Liu Y, Zhou R. 2017b. Comparative analysis of the cytology and transcriptomes of the cytoplasmic male sterility line H276A and its maintainer line H276B of cotton (Gossypium barbadense l.). INTERNATIONAL JOURNAL OF MOLECULAR SCIENCES 18(11): 2240.

Kühlbrandt W. 2015. Structure and function of mitochondrial membrane protein complexes. BMC BIOLOGY 13(1).

Levings CSII. 1993. Thoughts on Cytoplasmic Male Sterility in cms-T Maize. PLANT CELL 5(10): 1285–1290.

Levings CSPD. 1976. Restriction endonuclease analysis of mitochondrial DNA from normal and texas cytoplasmic Male-Sterile maize. SCIENCE 193(4248): 158–160.

Li M, Chen L, Khan A, Kong X, Khan MR, Rao MJ, Wang J, Wang L, Zhou R. 2021. Transcriptome and MiRNAomics Analyses Identify Genes Associated with Cytoplasmic Male Sterility in Cotton (Gossypium hirsutum L.). INTERNATIONAL JOURNAL OF MOLECULAR SCIENCES 22(9): 4684.

Ling Huang CZMF. 2011. The cell death in CMS plants. International Journal of Bioscience, Biochemistry and Bioinformatics 1(4): 297–301.

Liu Y, Zhou B, Khan A, Zheng J, Dawar FU, Akhtar K, Zhou R. 2021. Reactive oxygen species accumulation strongly allied with genetic male sterility convertible to cytoplasmic male sterility in kenaf. INTERNATIONAL JOURNAL OF MOLECULAR SCIENCES 22(3): 1107.

Livak KJ, Schmittgen TD. 2001. Analysis of relative gene expression data using Real-Time quantitative PCR and the 2^-ΔΔCT^ method. METHODS 25(4): 402–408.

Luo D, Xu H, Liu Z, Guo J, Li H, Chen L, Fang C, Zhang Q, Bai M, Yao N, Wu H, Wu H, Ji C, Zheng H, Chen Y, Ye S, Li X, Zhao X, Li R, Liu Y. 2013. A detrimental mitochondrial-nuclear interaction causes cytoplasmic male sterility in rice. NATURE GENETICS 45(5): 573–577.

Møller IM. 2001. PLANT MITOCHONDRIA and OXIDATIVE STRESS: Electron transport, NADPH turnover, and metabolism of reactive oxygen species. Annual Review of Plant Physiology and Plant Molecular Biology 52(1): 561–591.

Moneger F SCJL. 1994. Nuclear restoration of cytoplasmic male sterility in sunflower is associated with the tissue-specific regulation of a novel mitochondrial gene. EMBO 1(13): 8–17.

Narayanan S, Surolia A, Karande AA. 2004. Ribosome-inactivating protein and apoptosis: Abrin causes cell death via mitochondrial pathway in Jurkat cells. BIOCHEMICAL JOURNAL 377(1): 233–240.

Nie H, Cheng C, Hua J. 2020. Mitochondrial proteomic analysis reveals that proteins relate to oxidoreductase activity play a central role in pollen fertility in cotton. Journal of Proteomics 225: 103861.

Nigam M, Mishra AP, Salehi B, Kumar M, Sahrifi-Rad M, Coviello E, Iriti M, Sharifi-Rad J. 2019. Accelerated ageing induces physiological and biochemical changes in tomato seeds involving MAPK pathways. SCIENTIA HORTICULTURAE 248: 20–28.

Pfeffer S, Woellhaf MW, Herrmann JM, Förster F. 2015a. Organization of the mitochondrial translation machinery studied in situ by cryoelectron tomography. Nature Communications 6(1): 6019.

Pfeffer S, Woellhaf MW, Herrmann JM, Förster F. 2015b. Organization of the mitochondrial translation machinery studied in situ by cryoelectron tomography. Nature Communications 6(1): 6019.

Rhiannon R. White CLIL. 2020. Miro2 tethers the ER to mitochondria to promote mitochondrial fusion in tobacco leaf epidermal cells. communications biology.

Robles P, Quesada V. 2017. Emerging roles of mitochondrial ribosomal proteins in plant development. INTERNATIONAL JOURNAL OF MOLECULAR SCIENCES 18(12): 2595.

Savitski MM, Wilhelm M, Hahne H, Kuster B, Bantscheff M. 2015. A scalable approach for protein false discovery rate estimation in large proteomic data sets. Molecular &amp;amp; Cellular Proteomics 14(9): 2394.

Shahzad K, Zhang X, Guo L, Qi T, Tang H, Zhang M, Zhang B, Wang H, Qiao X, Feng J, Wu J, Xing C. 2020. Comparative transcriptome analysis of inbred lines and contrasting hybrids reveals overdominance mediate early biomass vigor in hybrid cotton. BMC GENOMICS 21(1).

Sies H, Jones DP. 2020. Reactive oxygen species (ROS) as pleiotropic physiological signalling agents. Nat Rev Mol Cell Biol 21(7): 363–383.

Sofo A, Dichio B, Xiloyannis C, Masia A. 2004. Effects of different irradiance levels on some antioxidant enzymes and on malondialdehyde content during rewatering in olive tree. Plant ence 166(2): 293–302.

Song J, Hedgcoth C. 1994. A chimeric gene (orf256) is expressed as protein only in cytoplasmic male-sterile lines of wheat. PLANT MOLECULAR BIOLOGY 26(1): 535–539.

Song S, Wang T, Li Y, Hu J, Kan R, Qiu M, Deng Y, Liu P, Zhang L, Dong H, Li C, Yu D, Li X, Yuan D, Yuan L, Li L. 2021. A novel strategy for creating a new system of third-generation hybrid rice technology using a cytoplasmic sterility gene and a genic male - sterile gene. PLANT BIOTECHNOLOGY JOURNAL 19(2): 251–260.

Sun Q, Hu C, Hu J, Li S, Zhu Y. 2009. Quantitative proteomic analysis of CMS-Related changes in honglian CMS rice anther. The Protein Journal 28(7): 341.

Suzuki H, Rodriguez-Uribe L, Xu J, Zhang J. 2013. Transcriptome analysis of cytoplasmic male sterility and restoration in CMS-D8 cotton. PLANT CELL REPORTS 32(10): 1531–1542.

Tomal A, Kwasniak-Owczarek M, Janska H. 2019. An update on mitochondrial ribosome biology: The plant mitoribosome in the spotlight. Cells 8(12): 1562.

Waltz F, Corre N, Hashem Y, Giegé P. 2020. Specificities of the plant mitochondrial translation apparatus. MITOCHONDRION 53: 30–37.

Wan C, Li S, Wen L, Kong J, Wang K, Zhu Y. 2007. Damage of oxidative stress on mitochondria during microspores development in Honglian CMS line of rice. PLANT CELL REPORTS 26(3): 373–382.

Wang Z, Qin G, Zhao TC. 2014. HDAC4: Mechanism of regulation and biological functions. Epigenomics 6(1): 139–150.

Wen B, Zhou R, Feng Q, Wang Q, Wang J, Liu S. 2014. IQuant: An automated pipeline for quantitative proteomics based upon isobaric tags. PROTEOMICS 14(20): 2280–2285.

Wolko J, Dobrzycka A, Bocianowski J, Bartkowiak-Broda I. 2019. Estimation of heterosis for yield-related traits for single cross and three-way cross hybrids of oilseed rape (Brassica napus L.). EUPHYTICA 215(10): 156.

Wu J, Zhang M, Zhang B, Zhang X, Guo L, Qi T, Wang H, Zhang J, Xing C. 2017. Genome-wide comparative transcriptome analysis of CMS-D2 and its maintainer and restorer lines in upland cotton. BMC GENOMICS 18(1).

Wu P, Xiao C, Cui J, Hao B, Zhang W, Yang Z, Ahammed GJ, Liu H, Cui H. 2020. Nitric oxide and its interaction with hydrogen peroxide enhance plant tolerance to low temperatures by improving the efficiency of the calvin cycle and the ascorbate – glutathione cycle in cucumber seedlings. JOURNAL OF PLANT GROWTH REGULATION.

Yang L, Wu Y, Zhang M, Zhang J, Stewart JM, Xing C, Wu J, Jin S. 2018a. Transcriptome, cytological and biochemical analysis of cytoplasmic male sterility and maintainer line in CMS-D8 cotton. PLANT MOLECULAR BIOLOGY 97(6): 537–551.

Yang L, Wu Y, Zhang M, Zhang J, Stewart JM, Xing C, Wu J, Jin S. 2018b. Transcriptome, cytological and biochemical analysis of cytoplasmic male sterility and maintainer line in CMS-D8 cotton. PLANT MOLECULAR BIOLOGY 97(6): 537–551.

Yang P, Han J, Huang J. 2014. Transcriptome sequencing and de novo analysis of cytoplasmic male sterility and maintenance in JA-CMS cotton. PLoS One 9(11): e112320.

Zhao H, Wang J, Qu Y, Peng R, Magwanga RO, Liu F, Huang J. 2020. Transcriptomic and proteomic analyses of a new cytoplasmic male sterile line with a wild Gossypium bickii genetic background. BMC GENOMICS 21(1).

Zheng J, Kong X, Li B, Khan A, Li Z, Liu Y, Kang H, Ullah Dawar F, Zhou R. 2019. Comparative transcriptome analysis between a novel allohexaploid cotton progeny CMS line LD6A and its maintainer line LD6B. INTERNATIONAL JOURNAL OF MOLECULAR SCIENCES 20(24): 6127.

Zhou B, Liu Y, Chen Z, Liu D, Wang Y, Zheng J, Liao X, Zhou AR. 2019. Comparative transcriptome analysis reveals the cause for accumulation of reactive oxygen species during pollen abortion in cytoplasmic Male-Sterile kenaf line 722HA. INTERNATIONAL JOURNAL OF MOLECULAR SCIENCES 20(21): 5515.

